# Lonafarnib improves cardiovascular function and survival in a mouse model of Hutchinson-Gilford Progeria Syndrome

**DOI:** 10.1101/2021.12.17.473197

**Authors:** S-I. Murtada, N. Mikush, M. Wang, P. Ren, Y. Kawamura, A.B. Ramachandra, D.T. Braddock, G. Tellides, L.B. Gordon, J.D. Humphrey

## Abstract

Clinical trials have demonstrated that lonafarnib, a farnesyltransferase inhibitor, extends lifespan in patients afflicted by Hutchinson-Gilford progeria syndrome, a devastating condition that accelerates many characteristics of aging and results in premature death due to cardiovascular sequelae. The US Food and Drug Administration approved Zokinvy™ (lonafarnib) in November 2020 for treating these patients, yet a detailed examination of drug-associated effects on cardiovascular structure, properties, and function has remained wanting. In this paper, we report encouraging outcomes of daily post-weaning treatment with lonafarnib on the composition and biomechanical phenotype of elastic and muscular arteries as well as associated cardiac function in a well-accepted mouse model of progeria that exhibits severe end-stage cardiovascular disease. Lonafarnib resulted in 100% survival of the treated progeria mice to the study end-point (time of 50% survival of untreated mice), with associated improvements in arterial structure and function working together to significantly reduce pulse wave velocity and improve left ventricular diastolic function. By contrast, dual treatment with lonafarnib and rapamycin did not improve outcomes over that achieved with lonafarnib monotherapy.

## INTRODUCTION

Hutchinson-Gilford progeria syndrome (HGPS) is an ultra-rare condition that results from a *de novo* autosomal dominant mutation to the gene (*LMNA*) that codes the nuclear envelope scaffolding protein lamin A (De Sandre-Giovannoli et al., 2003; Eriksson et al., 2003), which results in intranuclear accumulation of an aberrant form of the protein called progerin. This accumulation results from a variant that lacks a post-translational proteolytic processing step, leaving the progerin protein permanently modified by a farnesyl group. Clinical trials show that farnesyltransferase inhibitors extend healthspan and lifespan in children afflicted with HGPS (Gordon et al., 2012, 2014, 2018), and the US Food and Drug Administration approved on 20 November 2020 the use of the Zokinvy™ (lonafarnib) to slow the accumulation of progerin with the hope of extending lifespan (Dhillon, 2021).

Although HGPS is characterized by wide-spread tissue and organ abnormalities, premature death appears to result from cardiovascular complications, including atherosclerosis-related heart failure, with progressive left ventricular (LV) diastolic dysfunction the most prevalent condition observed in a recent clinical study (Prakash et al., 2018). It is well known in the normal aging population that increased aortic stiffness, and attendant increases in central pulse wave velocity (PWV), contributes significantly to both atherosclerosis and diastolic dysfunction / heart failure with preserved ejection fraction (Abhayaratna et al., 2006; van Popele et al., 2006; Desai et al., 2009; Townsend et al., 2015). Importantly, marked increases in ankle-brachial and carotid-femoral PWV have been reported in children with HGPS (Gerhard-Herman et al., 2012) and early results suggested that lonafarnib can lessen this increase (Gordon et al., 2012).

Notwithstanding the promise of lonafarnib in extending lifespan in HGPS patients, the largest and most recent study evaluated mortality, comparing treated trial patients with a contemporaneous untreated control group as its primary outcome (Gordon et al., 2018). That is, effects of lonafarnib on central artery stiffness and associated cardiovascular function have not been examined in detail. In this paper, we quantify and compare diverse functional cardiovascular metrics, including elastic and muscular artery stiffness and vasoactivity as well as cardiac performance, in untreated and lonafarnib-treated progeria mice (homozygous *Lmna*^*G609G/G609G*^ mice, denoted herein as G609G or simply GG). The original report of this mouse model indicated 50% survival at 242 days in the heterozygous (G609G/+) animals but only 103 days in the homozygous (G6096/G609G) animals (Osorio et al., 2011). All final assessments of structure and function presented herein for littermate wild-type (WT) and homozygous G609G mice are at 168/169 days of age, with improved survival of untreated progeria mice achieved via an improved feeding and care protocol that allowed the cardiovascular phenotype to worsen similar to that observed clinically in patients. Importantly, daily post-weaning treatment with lonafarnib improved cardiovascular function and yielded 100% survival to the study end-point. Dual treatment with lonafarnib and rapamycin, an inhibitor of mechanistic target of rapamycin (mTOR), did not improve outcomes over lonafarnib alone though these latter findings must be interpreted cautiously.

## RESULTS

### The aortic phenotype worsens rapidly in the perimorbid period in progeria

Switching from normal chow placed on the floor of the cage to a soft gel-based chow on the floor coupled with the introduction of a caretaker mouse in each cage extended the mean lifespan of untreated progeria mice by an additional ∼12%, from ∼150 days (Murtada et al., 2020) to ∼168 days, at which time cardiovascular function was evaluated. Figure 1A shows passive pressure-radius responses (reflecting the underlying microstructural composition and organization) of the descending thoracic aorta (DTA) in untreated progeria mice at multiple ages up to 168 days. There was a progressive structural stiffening (left-ward shift of the pressure-radius response) from 42 days to 140 days of age, with a remarkable stiffening thereafter to 168 days. This marked stiffening at 168 days associated with a grossly translucent, brittle appearance (Figure 1B), reflecting an extreme diffuse calcification.

**Figure 1.**
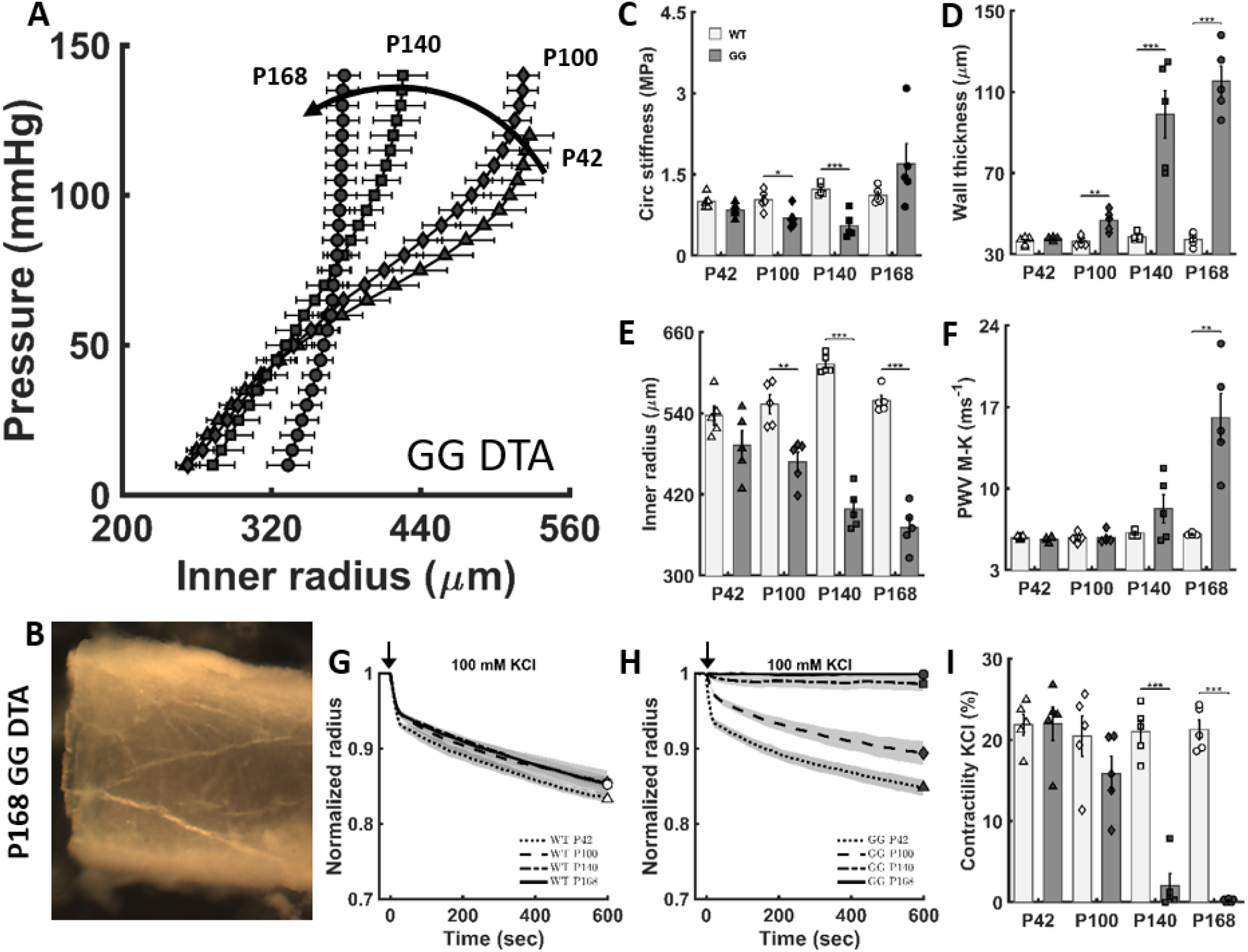
The biomechanical phenotype worsens progressively in the descending thoracic aorta (DTA) of untreated *Lmna*^*G6096G/G609G*^ progeria (GG) mice. (A) Pressure-diameter responses reveal a progressive structural stiffening as a function of age from 42 to 100 to 140 to 168 days (dark curved arrow). (B) Marked, late-stage aortic stiffening associated with a translucent appearance and brittle texture. (C-F) Passive geometric and mechanical metrics for the DTA in untreated progeria mice (dark bars) revealed marked progressive differences relative to wild-type (WT) littermates (light bars). (G-I) Vasoactive capacity, revealed as diameter reduction in response to high potassium depolarization of the smooth muscle cell membrane (100 mM KCl), first as a function of time and extent of response and second as steady state response, both at different ages. Note the near complete loss of contractility by 140 days and its complete loss by 168 days in progeria. *, **, and *** denote statistically significant differences between WT and GG at the age indicated, with *p* < 0.05, < 0.01, and < 0.001, respectively.

Quantification of material stiffness (Figure 1C) and aortic geometry (Figure 1D,E) enables calculation of the local aortic PWV via the Moens-Korteweg relation,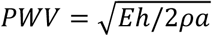, where *E* is wall stiffness, *h* wall thickness, *a* luminal radius, and *ρ* mass density of the blood. The circumferential material stiffness was lower in the G609G aorta at 42 days relative to that in WT littermate controls (0.68 MPa in G609G, 1.46 MPa in WT), but increased at 168 days (1.71 MPa in G609G, 1.86 MPa in WT). This late change, in combination with a marked increase in wall thickness (from 39 μm at 42 days to 115 μm at 168 days in G609G) and reduction in luminal radius (from 474 μm at 42 days to 371 μm at 168 days in G609G), increased the calculated local PWV (Figure 1F) by more than 3-fold in progeria at 168 days of age (from 5.3 m/s at 42 days to 16.2 m/s at 168 days). The vasoconstrictive capacity of this segment of the progeria aorta was also markedly abnormal (effectively absent) in the perimorbid period, with loss of contractile capacity seen by 140 days of age relative to control (Figure 1G-I). Figure S1A-G contrasts additional biomechanical metrics for the descending thoracic aorta from WT littermates and G609G mice, noting that biaxial wall stress and stretch were markedly diminished in progeria, and so too elastic energy storage capacity, especially in the perimorbid period. Because elastic energy storage during systolic distension normally facilitates the elastic recoil during diastole that augments blood flow, this marked loss of energy storage reveals a dramatic reduction in biomechanical function of the aorta in progeria.

### Lonafarnib improves survival

Only 53% (10/19) of the untreated progeria mice survived to postnatal day 168, the time of scheduled cardiac function assessment. By contrast, 100% (6/6) of the progeria mice treated daily with lonafarnib (450 mg/kg administered in the gel-based chow) from the time of weaning (at 21 days of age; Figure 2A) survived to complete cardiac function assessment at 168 days of age, noting that one of these 6 treated mice was found dead the day after anesthesia and echocardiography (Figure 2B). Three of the 10 untreated progeria mice that survived to day 168 and were not subjected to in vivo imaging were found dead on day 169, resulting in 7/19 (36.8%) untreated G609G mice surviving to day 169 versus 5/6 (83.3%) lonafarnib-treated G609G mice surviving to day 169. This increased survival of the treatment cohort emerged without a significant improvement in body mass (Figure 2C), with untreated and treated G609G mice having an ∼50% lower body mass than WT at 168 days of age.

**Figure 2.**
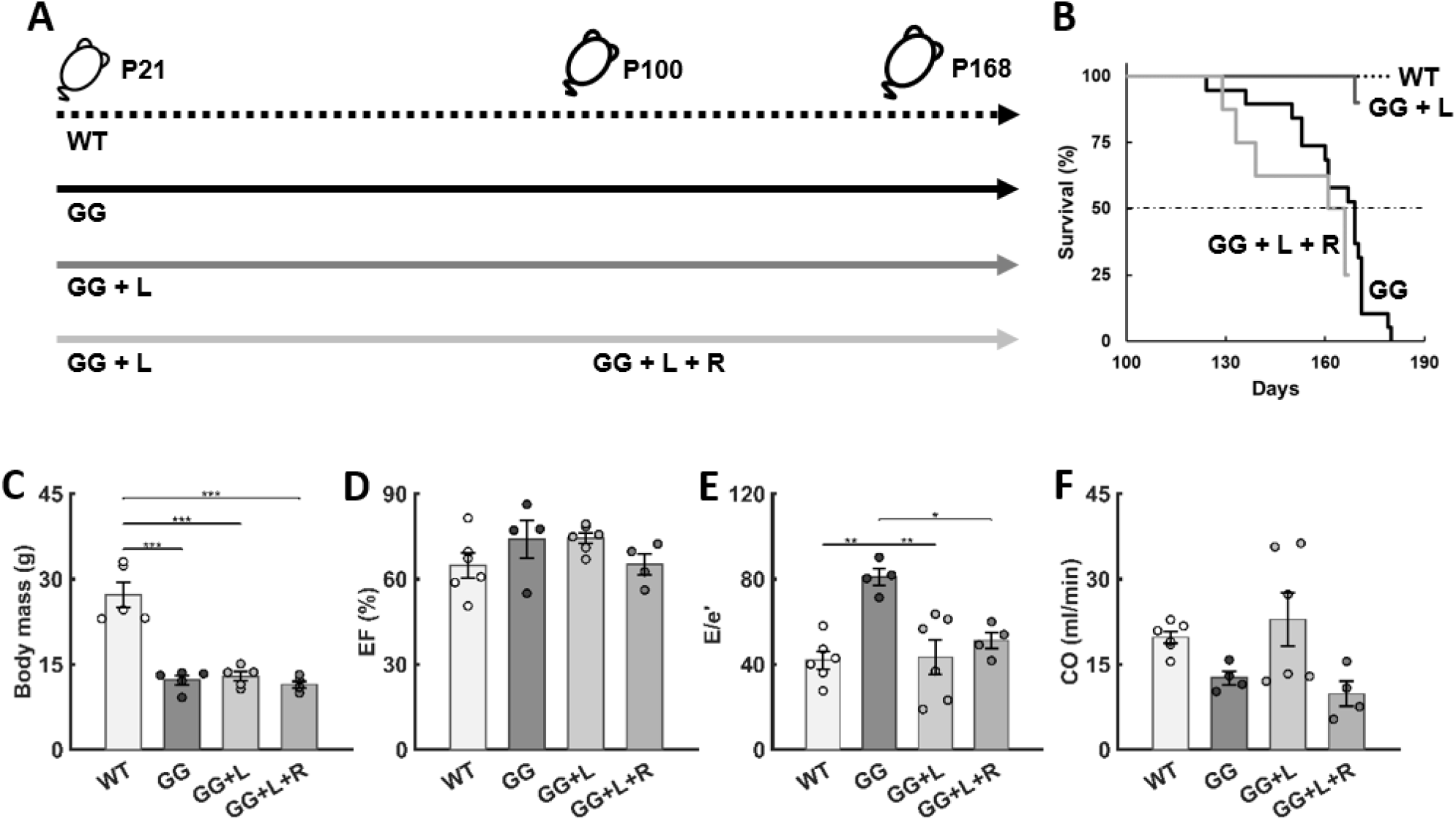
Lonafarnib improves survival and diastolic function. (A) Study design, including untreated wild-type (WT) littermate controls, untreated progeria (GG) mice, and progeria mice given lonafarnib (GG+L) daily in the chow from postnatal day P21 to P168. (B) All 6 lonafarnib-treated progeria mice (GG+L) survived to the intended end-point, P168, while 10 of the 19 untreated GG mice (∼53%) died by P169; note that the one GG+L death occurred at P169, the day after anesthesia and echocardiography. (C) Although lonafarnib did not improve body mass, it improved diastolic function (*E*/*e*’ in panel E) and cardiac output (*CO* in panel F) with no change in ejection fraction (*EF* in panel D). See Table 1 for all in vivo cardiovascular measurements. Finally, also shown are results from a combination therapy in progeria mice (GG+L+R), with lonafarnib given from P21 to P168 and rapamycin (R) given via IP injections from P100 to P168. There was no added benefit of the combination therapy, but instead a mortality similar to that in the untreated cohort. *, **, and *** denote statistical significance at *p* < 0.05, < 0.01, and < 0.001, respectively.

### Lonafarnib improves LV diastolic function and aortic PWV

Table 1 contrasts all cardiac measurements at 168 days of age for the three primary study groups: WT, untreated G609G, and lonafarnib-treated G609G. Most dimensioned cardiac metrics, such as LV chamber volumes and stroke volume (SV), were lower in both untreated and lonafarnib-treated progeria mice as expected allometrically based on their lower body mass (Murtada et al., 2020). Non-dimensioned metrics such as fractional shortening (FS) and ejection fraction (EF) remained similar across all three groups (WT, untreated and treated G609G mice) at 168 days of age (Figure 2D), indicating no loss in systolic function (and by inference, no change with treatment). Importantly, a key measure of diastolic function, related to LV filling velocity and mitral motions, improved in the progeria mice with lonafarnib treatment (Figure 2E) consistent with a trend toward an improved cardiac output (Figure 2F), which may explain, in part, their increased survival.

**Table 1.** Comparison of key cardiovascular metrics among wild-type (WT) mice and both untreated (GG) and lonafarnib treated (GG+L) progeria mice. An * denotes a statistically significant difference at *p* < 0.05, and a ** denotes significance at *p* < 0.01, both between untreated and treated GG. IVS – interventricular septum, LW – left ventricular free wall, CO – cardiac output, EF – ejection fraction, FS – fractional shortening, LV – left ventricular, E – early and A – late ventricular filling blood velocities, the latter due to atrial contraction, e’ – early and a’ – late tissue velocities, and DT – deceleration time. A lower E/A ratio indicates diastolic dysfunction. The * and ** denote statistical differences at *p* < 0.05 or 0.01 between treated (either GG+L or GG+L+R) and untreated (GG) progeria mice, the key comparisons of interest.

|  | WT | GG | GG + L | GG + L + R |
| --- | --- | --- | --- | --- |
| *n* | 6 | 4 | 6 | 4 |
| *Heart rate (bpm)* | 431 ± 20 | 405 ± 49 | 426 ± 32 | 411 ± 28 |
| *Diastolic thickness (mm)* |  |  |  |  |
| IVS | 0.73 ± 0.03 | 0.70 ± 0.03 | 0.75 ± 0.06 | 0.68 ± 0.12 |
| LW | 0.67 ± 0.06 | 0.69 ± 0.05 | 0.78 ± 0.02 | 0.59 ± 0.09 |
| *Internal Diameter (mm)* |  |  |  |  |
| Diastolic | 4.04 ± 0.07 | 3.35 ± 0.17 | 3.30 ± 0.06 | 3.20 ± 0.04 |
| Systolic | 2.61 ± 0.18 | 1.93 ± 0.26 | 1.90 ± 0.07 | 2.08 ± 0.03 |
| *Volume (ul)* |  |  |  |  |
| Diastolic | 72 ± 3 | 46 ± 5 | 44 ± 2 | 42 ± 6 |
| Systolic | 26 ± 4 | 13 ± 4 | 11 ± 1 | 14 ± 2 |
| SV (ul) | 46 ± 1 | 34 ± 3 | 33 ± 1 | 28 ± 5 |
| *CO (mL/min)* | 20 ± 1 | 13 ± 1 | 23 ± 5 | 10 ± 2 |
| *EF (%)* | 65 ± 4 | 74 ± 7 | 74 ± 2 | 65 ± 4 |
| *FS (%)* | 36 ± 3 | 43 ± 6 | 42 ± 2 | 35 ± 3 |
| *LV Mass (mg)* | 81 ± 5 | 59 ± 8 | 65 ± 4 | 48 ± 5 |
| *LV/Body Mass (mg/g)* | 3.2 ± 0.3 | 4.3 ± 0.3 | 5.3 ± 0.3* | 4.0 ± 0.5 |
| *Filling Velocity (mm/s)* |  |  |  |  |
| E Peak | 666 ± 27 | 620 ± 99 | 566 ± 59 | 458 ± 83 |
| A Peak | 384 ± 59 | 377 ± 46 | 300 ± 42 | 222 ± 44 |
| Septal e' | 19.08 ± 1.53 | 7.34 ± 1.40 | 12.91 ± 1.43* | 7.08 ± 1.11 |
| Septal a' | 21.32 ± 0.94 | 12.34 ± 1.87 | 11.6 ± 0.6 | 9.1 ± 0.2 |
| DT (ms) | 26.4 ± 1.6 | 31.6 ± 4.2 | 25.5 ± 2.3 | 23.0 ± 2.8 |
| *Velocity Ratios (-)* |  |  |  |  |
| E/A | 2.03 ± 0.42 | 1.63 ± 0.12 | 1.95 ± 0.12 | 2.79 ± 1.31 |
| Septal e'/a' | 0.91 ± 0.11 | 0.60 ± 0.07 | 1.14 ± 0.15* | 0.79 ± 0.14 |
| Average E/e' | 41.99 ± 4.21 | 80.97 ± 3.86 | 43.41 ± 8.06** | 51.21 ± 3.85** |

Figure 3 shows results similar to those in Figure 1 except for the effects of lonafarnib on aortic function. Note the passive pressure-radius behavior of the descending thoracic aorta at 168 days of age, contrasting that for WT with that of untreated and lonafarnib-treated G609G mice (panel A). The structural stiffening in progeria slightly was less with treatment, as evidenced by a right-ward shift of the pressure-radius response. There were also modest improvements in all three contributors to PWV, namely, circumferential material stiffness (panel B), wall thickness (panel C), and luminal radius (panel D). Importantly, these three changes worked together to improve aortic PWV significantly, lowering the calculated value by 36.5%, from 16.2 m/s for the untreated progeria mice to 10.3 m/s in the treated mice.

**Figure 3.**
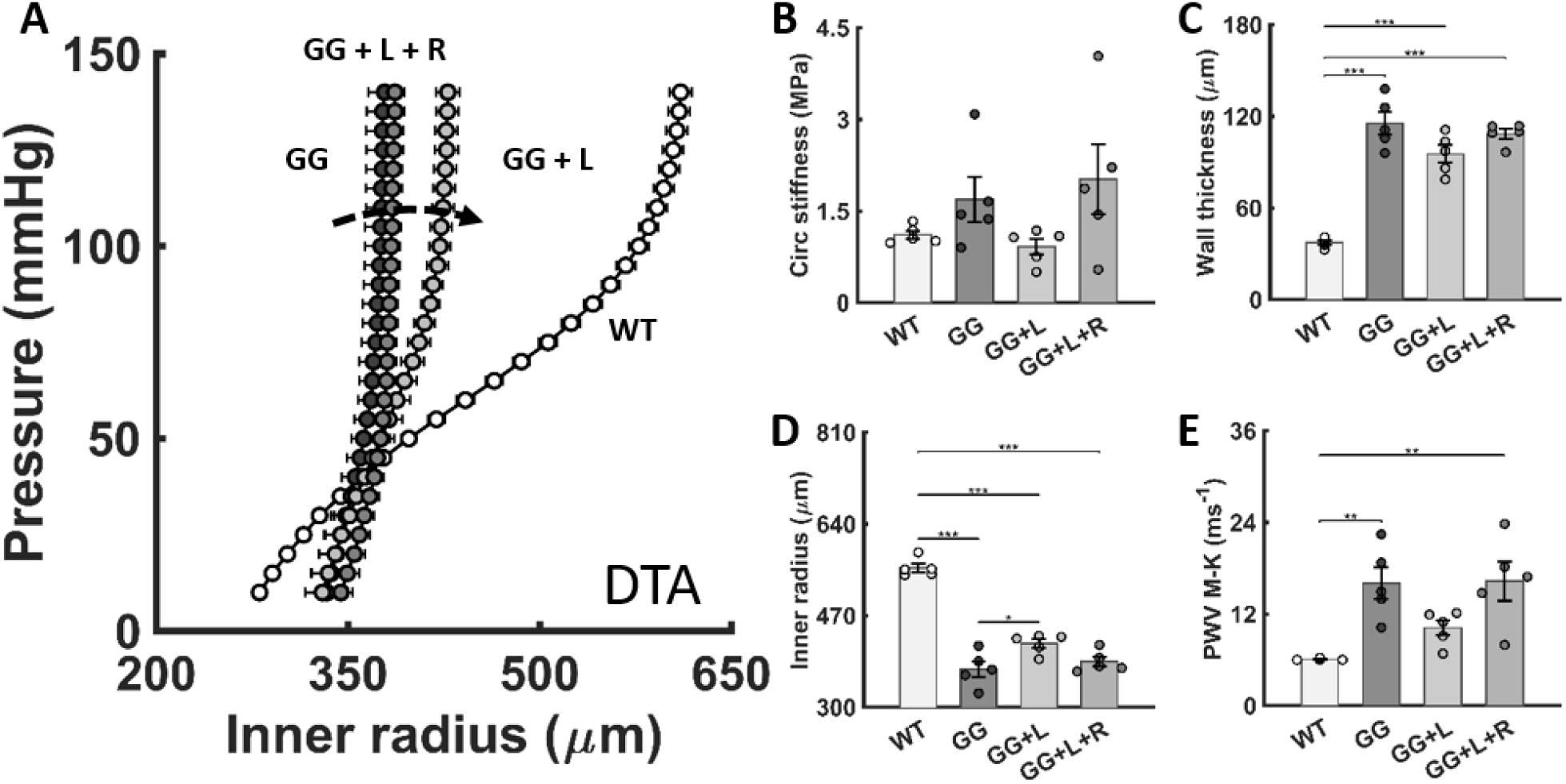
Lonafarnib improves PWV at 168 days of age via modest but complementary changes in aortic geometry and properties. (A) Pressure-diameter results comparing untreated (GG) and lonafarnib treated (GG+L) descending thoracic aortas (DTA) versus those from wild-type (WT). (B-E) Modest (non-significant) decreases in circumferential material stiffness and wall thickness and increases in inner radius yet worked together to yield a significant reduction in PWV with lonafarnib treatment. Shown, too, are results from a study combining lonafarnib and rapamycin treatment (GG+L+R), which revealed no further benefit. The dashed black arrow indicates changes with lonafarnib treatment; *, **, and *** denote statistical significance at *p* < 0.05, < 0.01, and < 0.001, respectively.

### Lonafarnib improves aortic composition

Histological sections of the descending thoracic aorta at 168/169 days of age for the WT littermate controls and untreated and lonafarnib-treated G609G mice (Figure 4A) revealed that lonafarnib treatment resulted in a significantly lower accumulation of mural proteoglycans in progeria (Figure 4B) and less mural calcification that manifested between 140 and 168 days in progeria (Figure 4C). Both of these changes likely contributed to the improved PWV, in part via a reduced wall thickness (Table S1). This improved PWV emerged despite no improvement with lonafarnib treatment in either the elastic energy storage (Table S1) or the vasoconstrictive capacity of this segment of the aorta (Table S2). All other passive and active geometric and biomechanical metrics are similarly tabulated in Tables S1 and S2.

**Figure 4.**
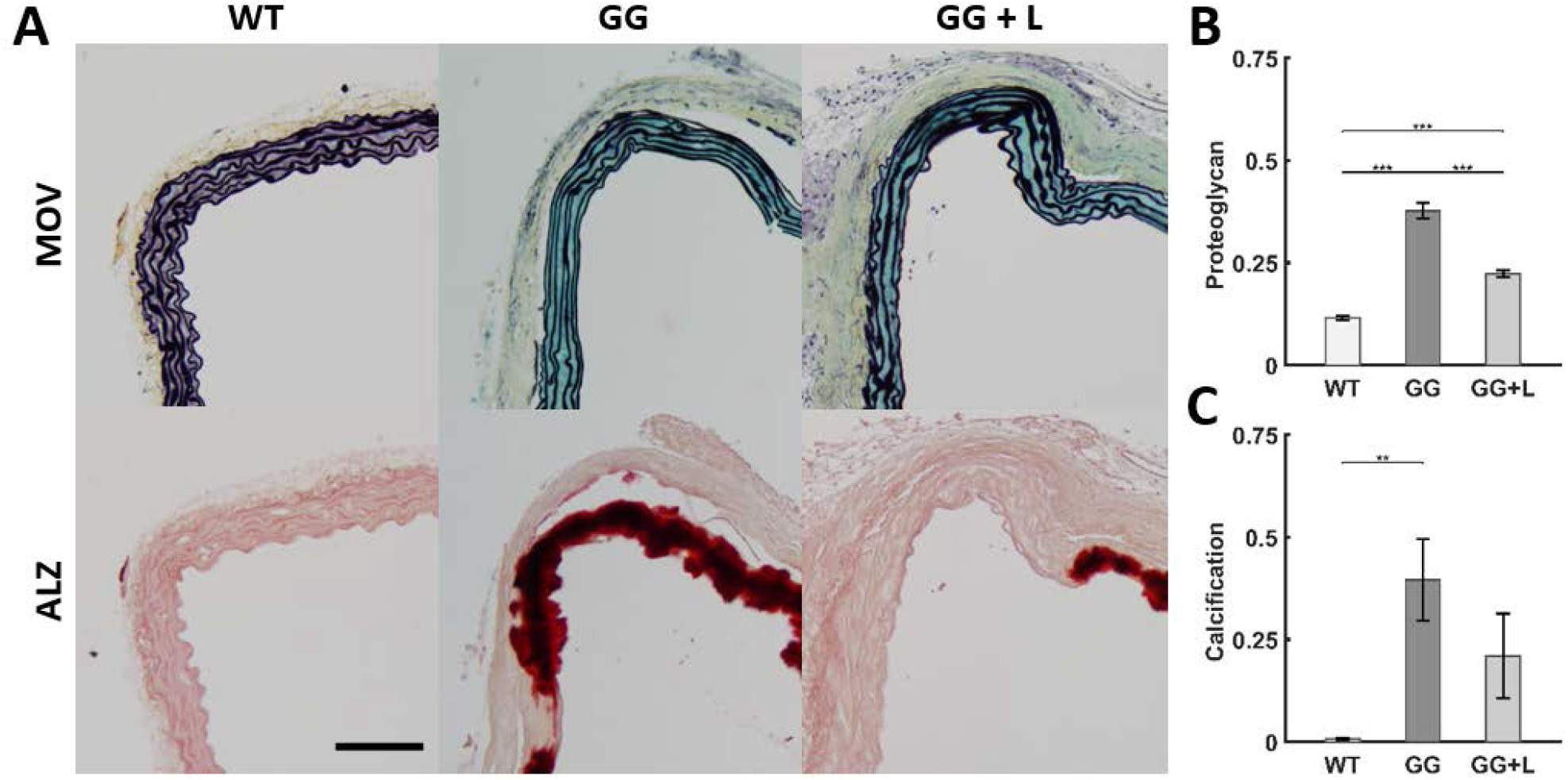
Lonafarnib treatment reduces proteoglycan accumulation and calcification at 168 days of age within the descending thoracic aorta (DTA) in progeria mice. A (First row) Movat-stained sections reveal intact elastic laminae (dark lines) but excessive accumulations of proteoglycans (blue staining) in the media and adventitia at post-natal day 168 in progeria (GG) relative to littermate wild-type (WT) controls; (B) this accumulation is reduced significantly in progeria following lonafarnib treatment (GG+L). A (Second row) Alizarin red-stained sections reveal marked calcification of portions of the DTA in progeria, (C) which was also reduced in some cases with lonafarnib treatment. *, **, and *** denote statistical significance at *p* < 0.05, < 0.01, and < 0.001, respectively. The scale bar is 100 microns.

### Lonafarnib differentially affects elastic and muscular arteries

Elastic (e.g., the aorta) and muscular (e.g., branch mesenteric) arteries remodel differently in hypertension and natural aging (Laurent and Boutouyrie, 2015; Murtada et al., 2021). We thus quantified the biomechanical phenotype of the second order mesenteric artery (MA) as a representative muscular artery. Both the structural (pressure-radius response; Figure 5A) and material (Figure 5B) stiffness were compromised by progeria despite no apparent histological signs of calcification or excessive accumulation of proteoglycans (Figure S2). Wall thickness was increased (Figure 5C), but luminal radius decreased (Figure 5D), thus resulting in no change in calculated local PWV (Figure 5E). Most dramatic, however, was a significant decrease in vasocontractility at 168 days of age in the progeria mice (Figure 5F-I). Treatment with lonafarnib resulted in a modest leftward shift in the passive pressure-radius behavior relative to untreated but without significant effects on passive geometry and properties (Figure 5A-E). Importantly, however, lonafarnib largely prevented the decline in vasoconstrictive and vasodilatory capacity seen in progeria vessels that were untreated (Figure 5F-I). This effect is expected to result in an increased capacity and ability to regulate lumen size and blood flow in this muscular artery.

**Figure 5.**
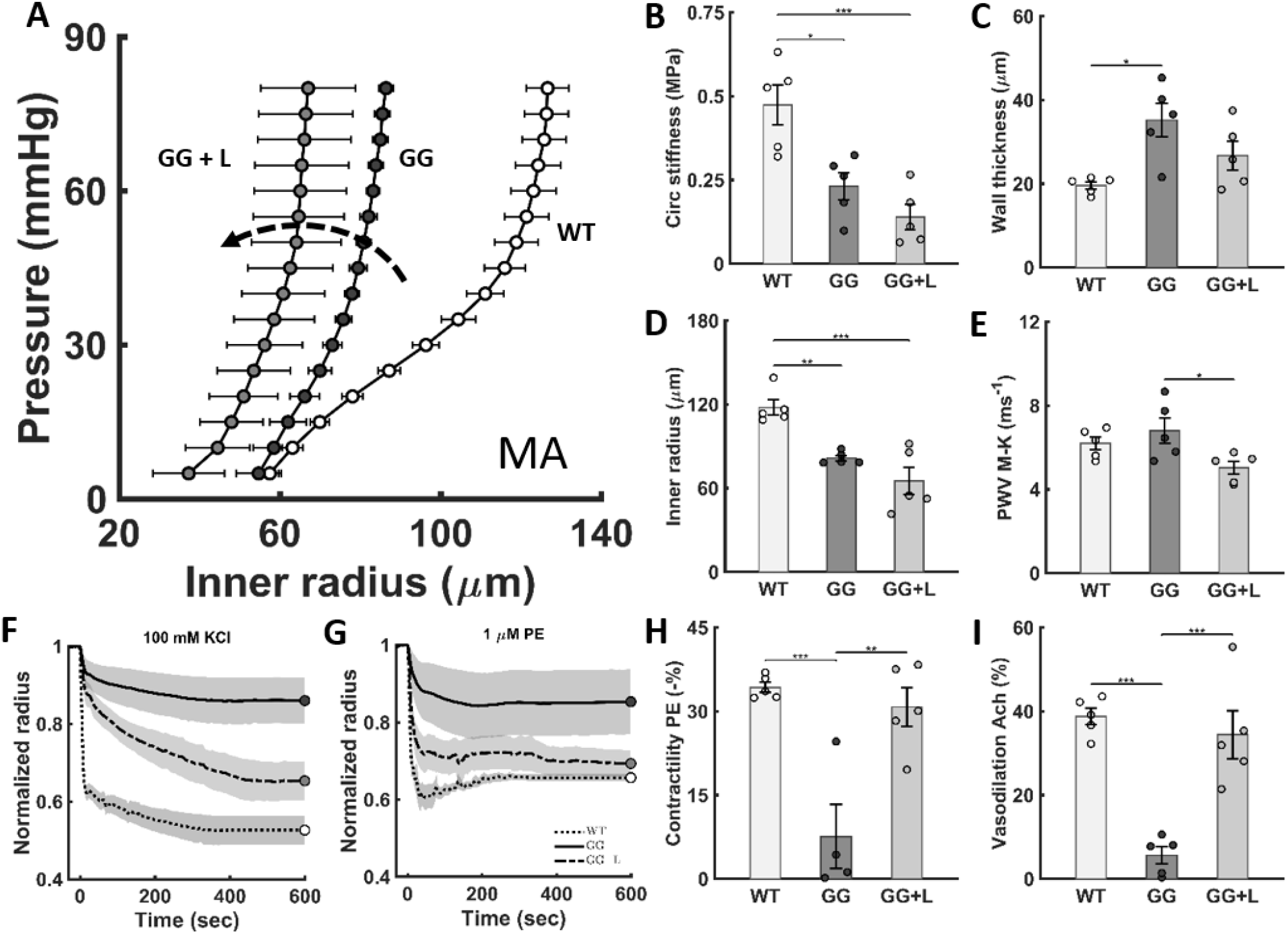
Effects of lonafarnib on geometric and mechanical metrics for the second branch mesenteric artery (MA). (A) Progeria compromises the structural stiffness of the mesenteric artery at 168 days of age, and lonafarnib increases this structural stiffening further. (B-E) Progeria compromises the circumferential material stiffness, wall thickness, and inner radius of the mesenteric artery, though resulting in modest changes in calculated PWV, with lonafarnib reducing this value. (F-I) Progeria compromises the vasocontractile capacity of the mesenteric artery, which is rescued by lonafarnib. Note that endothelial cell function, as indicated by dilatation of pre-constricted vessels with acetylcholine, appears not to be affected by progeria or its treatment. The dashed black arrow indicates changes with lonafarnib treatment; the *, **, and *** denote statistical significance at *p* < 0.05, < 0.01, and < 0.001, respectively.

### Combined lonafarnib and rapamycin did not improve outcomes further

Finally, we examined possible benefits afforded by a combination therapy, daily intraperitoneal injections of the mTOR inhibitor rapamycin from day 100 to day 168 in progeria mice that received lonafarnib daily from day 21 to day 168 via the soft chow (Figure 2A). As seen in Figure 2, there was no improvement in weight gain (panel C) or diastolic function (panel E), but rather a worse survival (panel B) and reduced cardiac output (panel F) relative to lonafarnib treatment alone despite no effect on ejection fraction (panel C). There were similarly no benefits to large artery function (Figure 3A-D), but rather a slight worsening of calculated PWV in the descending thoracic aorta (Figure 3E). Effects of combination treatment were not assessed in the mesenteric artery, which presented with a milder progeria phenotype.

## DISCUSSION

Prior studies have documented marked changes in arterial structure in HGPS, including loss of vascular smooth muscle cells, accumulation of proteoglycans, and increased fibrillar collagen, both in patients (Stehbens et al., 2001; Olive et al., 2010) and in mouse models (Varga et al., 2006; Capell et al., 2008; Kim et al., 2018). Vascular calcification has also been reported in patients and aged heterozygous G609G mice (Villa-Bellosta et al., 2013) as well as in homozygous G609G mice crossed with *Apoe*^-/-^ mice (Hamczyk et al., 2018). We recently showed in homozygous G609G mice that the marked intramural accumulation of proteoglycans in the aortic wall was a driving factor for increasing PWV in the absence of calcification at 140 days of age (Murtada et al., 2020). Here, we demonstrate a further rapid increase in PWV in the perimorbid period in these mice (∼16 m/s at day 168), driven in part by extensive aortic medial calcification as observed in some patients. Furthermore, we found a continued loss of smooth muscle contractile capacity at 168 days, consistent with findings at 140 days of age (Murtada et al., 2020) as well as with reports of reduced responsiveness to vasodilators (Varga et al., 2006).

It has been suggested that G609G mice die from starvation and cachexia, with demonstration that a high-fat diet can improve survival in these mice, ∼74% from a mean of 111 days to 193 days (Kreienkamp et al., 2018). Given the potential role of atherosclerosis in the cardiovascular pathology, however, we used a normal gel-based diet that facilitated consumption and helped to improve survival to a maximum of ∼180 days, with 53% survival at 168 days. It is noted that this gel-based diet is a nutritionally fortified dietary supplement that combines hydration and nutrition, but with equivalent levels of fat as regular chow. We submit that this increased survival without the added complications of a high-fat diet allowed time for a severe cardiovascular phenotype that more closely mimics that reported in HGPS patients to manifest. Thus, we were able to study the effects of drug intervention on certain key characteristics of severe cardiovascular disease for the first time.

HGPS is a mechano-sensitive condition (Kirby and Lammerding, 2018), with stiff tissues and organs most affected. The normal murine aorta is highly stressed by hemodynamic loads and it normally expresses high levels of lamin A (Kim et al., 2018). Consistent with this observation, we found the progeroid phenotype to be much more severe in the descending thoracic aorta (with passive intramural stresses > 200 kPa in normalcy) than in the mesenteric artery (with stresses < 65 kPa in normalcy). Reduced contractile responses were marked, however, in both the aorta and mesenteric artery. Lonafarnib treatment resulted in only modest long-term improvements in passive stiffness and geometry of the aorta, yet these effects worked together to reduce PWV significantly likely due to the reduction in mural proteoglycans and calcification despite not rescuing the contractile phenotype. It was previously shown that high dose tipifarnib also reduced proteoglycan accumulation in the aorta (cf. Capell et al., 2008) though without detailed assessment of functional consequences. By contrast, lonafarnib had modest effects on the passive properties of the mesenteric artery but largely preserved its vasoconstrictive and hence vasodilatory potential. This improved contractile capacity appears to have led to the modest reduction in caliber. It is well known that arterioles in the systemic vasculature tend to inward remodel in hypertension (due in part to their myogenic capacity), which may help prevent elevated pressure pulses from propagating into the microcirculation of end organs, thus protecting these organs (Boutouyrie et al., 2021). Indeed, we previously found a similar inward remodeling with increased vasoconstrictive capacity in the mesenteric artery in a mouse model of induced hypertension (Murtada et al., 2021). Although we did not assess coronary arteries or the coronary microcirculation, an increased vasoregulatory capacity of muscular arteries due to lonafarnib treatment could have combined with the improved central hemodynamics to improve LV diastolic function. In line with this, recent studies have suggested that heart failure with preserved ejection fraction, which is characterized by dysfunctional filling of the left ventricle during diastole, associates with coronary artery disease (Rush et al., 2021). There is, therefore, a need to investigate further the potential differential effects of lonafarnib on arteries within the many different vascular beds, particularly in end organs, noting again that the normal levels of hemodynamically-induced stresses differ along the vascular tree.

Although lonafarnib significantly increases lifespan in children with HGPS, disease remains progressive and patients still die early. Hence, multiple studies have considered combination or other therapies. One trial concluded, however, that there was no further cardiovascular benefit of adding the prenylation inhibitors pravastatin and zoledronic acid (a bisphosphonate) to lonafarnib therapy (Gordon et al., 2016). Improvement in survival was similarly unremarkable for these combination therapies in homozygous G608G mice despite improvements in the musculoskeletal phenotype (Cubria et al., 2020). By contrast, inhibitors of mTOR, namely rapamycin (Cao et al., 2011) and everolimus (DuBose et al., 2018), have shown some promise in vitro in treating progeria cells, and genetic reduction of mTOR in homozygous G608G progeria mice extended lifespan ∼30% (Cabral et al., 2021). We compared lonafarnib + rapamycin versus lonafarnib alone. Rapamycin was given from 100 to 168 days of age via daily intraperitoneal (IP) injections at 2 mg/kg, consistent with a dosing-regime shown to be effective in the treatment of aortopathies in mice (Li et al., 2014). We did not find any benefit of the combined therapy when assessing large artery structure or function, but instead found early mortality similar to that in untreated progeria mice. Given the frailty of these mice, one cannot exclude a detrimental consequence of daily handling and IP injections leading to increased emotional and physical stress, though G609G mice have been reported to tolerate two IP injections per week for 5 weeks (Lee et al., 2016). Without any clear benefit, our lonafarnib + rapamycin study was terminated though other methods of drug administration and other concentrations should be considered, particularly given the aforementioned promising results of genetic reductions in mTOR signaling, though germline manipulations also affect tissues developmentally.

Efforts by others are underway to use genetic manipulations to rescue the progeria phenotype, with one study reporting a 26% increase (Santiago-Fernandez et al., 2019) and another a 51% increase (Beyret et al., 2019) in median survival rate via disruption of *Lmna* and progerin transcription. Although these studies provide excellent proof-of-principle, genetic editing occurred mainly in the liver, not in the cardiovascular system, and there was no detailed assessment of potential changes in cardiovascular structure or function. Over 90% of children afflicted with HGPS exhibit the c.1824C>T, p.G608G mutation. Targeted anti-sense therapies in heterozygous and homozygous G608G mice have extended lifespan 33 to 62%, from ∼220 days to ∼330 days in these mice (Erdos et al., 2021; Pattaraju et al., 2021), which have a less severe phenotype than homozygous G609G mice. Most dramatically, gene editing in the homozygous G608G mouse extended lifespan more than 2-fold, from 215 to 510 days of age (Koblan et al., 2021). There is, therefore, significant promise for both genetic editing and RNA-based treatment strategies.

Nevertheless, until HGPS clinical trials establish safety and efficacy for genetics-based treatments, pharmacotherapy will likely remain a mainstay in patient care. Importantly herein, daily post-weaning treatment with lonafarnib yielded 100% survival of G609G mice to the scheduled time of anesthesia and echocardiography at 168 days of age, which can be contrasted with prior reports of extended 50% survival in these same G609G mice to 133 days (with progerinin treatment; Kang et al., 2021), 140 days (with JH4 treatment; Lee et al., 2016), 161 days (with genetic manipulation; Santiago et al., 2019), and 177 days (with genetic manipulation; Beyret et al., 2019). Recalling that LV diastolic dysfunction was the most prevalent cardiovascular abnormality observed in a recent clinical assessment of HGPS patients (Prakash et al., 2018), we submit that the increased survival in our *Lmna*^*G609G/G609G*^ mice resulting from daily lonafarnib monotherapy likely resulted from a combination of multiple modest improvements in large artery composition and function that worked together to significantly improve both central artery hemodynamics (PWV) and LV diastolic function, with improved small artery function also likely contributing by attenuating the propagation of pulse pressure waves into end organs. Importantly, these *Lmna*^*G609G/G609G*^ mice, which with lifespan extension via improved feeding and care, otherwise exhibited a particularly severe aortic phenotype in the perimorbid period. The present findings thus confirm promise and encourage use of lonafarnib clinically as the field continues to work to develop a definitive cure (progeriaresearch.org).

## METHODS

### Animals

All live animal procedures were approved by the Yale University Institutional Animal Care and Use Committee (IACUC). *Lmna*^*G609G/G609G*^ mice were originally developed by Osorio et al. (2011). Herein, mice were generated by breeding *Lmna*^*G609G/+*^ mice to obtain homozygous wild-type (*Lmna*^*+/+*^) and progeria (*Lmna*^*G609G/G609G*^, denoted herein as G609G or simply GG) mice, all having a C57BL/6 genetic background (Murtada et al., 2020). A total of 26 mixed-sex WT mice and 68 G609G mice were scheduled for study at 168 days of age. Following in vivo collection of cardiac data under anesthesia, the mice were euthanized using an intraperitoneal injection of Beuthanasia-D either the same (168) or next (169) day for tissue harvest, with death confirmed following exsanguination upon removal of the descending thoracic aorta.

### Drug Treatment

Lonafarnib was obtained from The Progeria Research Foundation Cell and Tissue Bank and given daily, at a dose of 450 mg/kg, within the soft gel-based chow (Clear H_2_O, DietGel 76A) from postnatal day 21 to the study end-point of 168/169 days of age. Rapamycin (Millipore Sigma), at a dose of 2 mg/kg, was given daily via intraperitoneal (IP) injections from 100 days to the study end-point of 168/169 days of age in the lonafarnib-treated cohort.

### Cardiac Function Testing

Following standard procedures (Ferruzzi et al., 2020; Murtada et al., 2020), mice were anesthetized with isoflurane at 168 days of age and data were collected using a Vevo 2100 ultrasound system (Visualsonic, Toronto, Canada) using a linear array probe (MS550D, 22-55 MHz). Systolic and diastolic function of the left ventricle (LV) were quantified using B-Mode imaging to provide parasternal long axis (LAX) and short axis (SAX) views, with M-Mode imaging in both planes to track the temporal evolution of cavity diameter at the level of the papillary muscles to measure LV systolic function. B-Mode images of the LV outflow tract diameter and pulsed-wave Doppler images of blood velocity patterns across the aortic valve estimated aortic valve area as a check for valve stenosis. LV diastolic function was monitored using an apical four-chamber view, which was achieved by aligning the ultrasound beam with the cardiac long axis. Color Doppler imaging located the mitral valve, while pulsed-wave Doppler measured mitral inflow velocity. Finally, Doppler tissue imaging from the lateral wall and interventricular septum measured the velocities of tissue motion.

Echocardiographic data were processed using Visualsonic data analysis software and standard methods. Global parameters of LV function were measured in triplicates from individual parasternal LAX and SAX views, and final parameters were obtained as averages from both views. Herein, LV systolic function is characterized by parameters such as cardiac output (CO) and ejection fraction (EF), while LV diastolic functional parameters include peak early (E) filling velocity and peak atrial (A) filling velocity, and their ratio (E/A), as well as the E-wave deceleration time (DT).

### Biomechanical Testing

Our biomechanical testing and analysis follows validated protocols (Ferruzzi et al., 2013; Murtada et al., 2020). Briefly, excised vessels (descending thoracic aorta and the second branch mesenteric artery) were mounted on custom-drawn glass cannulae and secured with 6-0 suture. They were then placed within a custom computer-controlled biaxial device for biomechanical testing and immersed in a heated (37°C) and oxygenated (95% O_2_, 5% CO_2_) Krebs-Ringer bicarbonate buffered solution containing 2.5 mM CaCl_2_. The vessels were then subjected to a series of isobaric (distending pressure of 90 mmHg for the aorta, 60 mmHg for the mesenteric artery) - axially isometric (fixed specimen-specific in vivo axial stretch) protocols wherein they were contracted with 100 mM KCl, relaxed (KCl washed out), contracted with 1 μM phenylephrine (PE), and then dilated with 10 µM acetylcholine (ACh; without wash-out), an endothelial cell dependent stimulant of nitric oxide synthesis.

Upon completion of these active protocols, the normal Krebs solution was washed out and replaced with a Ca^2+^-free Krebs solution to ensure a sustained passive behavior while maintaining heating and oxygenation. Vessels were then preconditioned via four cycles of pressurization (between 10 and 140 mmHg for the aorta, and between 10 and 90 mmHg for the mesenteric artery) while held fixed at their individual in vivo axial stretch. Finally, specimens were exposed to seven cyclic protocols: three pressure-diameter (*P-d*) protocols, with luminal pressure cycled between 10 and either 140 mmHg (aorta) or 90 mmHg (mesenteric) while axial stretch was maintained fixed at either the specimen-specific in vivo value or ±5% of this value, plus four axial force-length (*f-l*) tests, with force cycled between 0 and a force equal to the maximum value measured during the pressurization test at 5% above the in vivo axial stretch value, while the luminal pressure was maintained fixed at either {10, 60, 100, or 140 mmHg} for the aorta or {10, 30, 60, 90} for the mesenteric artery. Pressure, axial force, outer diameter, and axial length were recorded on-line for all seven of these protocols and used for subsequent data analysis (using the last unloading curve of each, which provides information on elastically stored energy that is available to work on the distending fluid), noting that prior studies have revealed robust fixed-point estimates of constitutive parameters based on such data.

### Passive Descriptors

Five key biomechanical metrics define well the passive mechanical phenotype of the aorta: mean circumferential and axial wall stress, circumferential and axial material stiffness linearized about a physiologic value of pressure and axial stretch, and the elastically stored energy. Each of these metrics is calculated easily given best-fit values of the model parameters within a nonlinear stored energy function, herein taken as (Ferruzzi et al., 2013; Murtada et al., 2020)

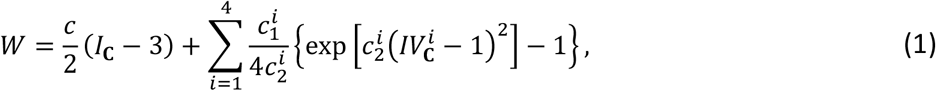

where *c* (dimension of kPa), 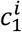 (dimension of kPa), and 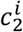 (dimensionless, with *i* = 1,2,3,4) are model parameters. *I*_**C**_ **= C: I** and 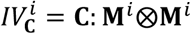 are coordinate invariant measures of deformation, with **I** the identity tensor and **C = F**^T^**F** the right Cauchy-Green tensor, with **F** the deformation gradient tensor (mapping from the traction-free configuration to any loaded configuration, which is pressurized and axially stretched herein) and superscript T the transpose; det **F** = 1 because of assumed incompressibility. The direction of the *i*^th^ family of fibers is given by the unit vector 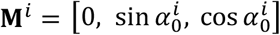, with angle 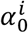 computed with respect to the axial direction in the traction-free reference configuration. Based on prior microstructural observations, and the yet unquantified effects of cross-links and physical entanglements amongst the multiple families of fibers, we included contributions of axial 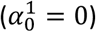, circumferential 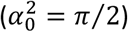, and two symmetric diagonal families of fibers 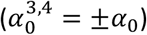 to capture phenomenologically the complex biaxial material behavior; this relation has been validated independently.

Theoretical values of the applied loads were computed from components of Cauchy stress by solving global equilibrium equations in the radial and axial directions. Best-fit values of the eight model parameters were estimated via nonlinear regression (Marquardt-Levenberg) to minimize the sum-of-the-squared differences between experimentally-measured and theoretically-predicted values of luminal pressure and axial force, each normalized by average experimental measures. Estimated parameters (constrained to be non-negative) were used to compute stress, material stiffness, and stored energy at any configuration. For example, components of the stiffness tensor (C_*ijkl*_), linearized about a configuration defined by the distending pressure and in vivo value of axial stretch, were computed as

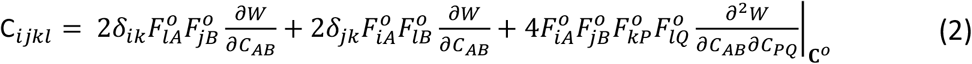

where *δ*_*ij*_ are components of **I, F**^*o*^ is the deformation gradient tensor between the chosen reference configuration and a finitely deformed in vivo relevant configuration, and **C**^*o*^ is the corresponding right Cauchy-Green tensor. Local aortic PWV was calculated using both the Moens-Korteweg (based on material stiffness and wall geometry) and Bramwell-Hill (based on overall wall compliance) equations, which yielded similar results. The former was preferred for it better delineates contributors to this important metric.

### Histology

Following biomechanical testing, specimens were unloaded and fixed overnight in 10% neutral buffered formalin, then stored in 70% ethanol at 4°C for histological examination. Fixed samples were dehydrated, embedded in paraffin, sectioned serially (5-μm thickness), and stained with Movat Pentachrome (showing elastic fibers in black, glycosaminoglycans in blue) or Alizarin Red (showing calcium in red) for standard histology. Detailed analyses were performed on three biomechanically representative vessels per group. Histological images were acquired on an Olympus BX/51 microscope (under bright light or fluorescence imaging) using an Olympus DP70 digital camera (CellSens Dimension) and a 20x magnification objective. Custom MATLAB scripts extracted layer-specific cross-sectional areas and calculated positively stained pixels and area fractions.

### Statistics

One- or two-way analysis of variance (ANOVA) with Bonferroni post-hoc testing was used to compare results, with *p* < 0.05 considered significant. All data are presented as mean ± standard error of the mean (SEM).

## ACKNOWLEDGMENTS

This work was supported, in part, by grants from the US National Institutes of Health (R01 HL105297, R21 AG067347) and Inozyme Pharma, Inc. (to D.T.B.). We also thank The Progeria Research Foundation (progeriaresearch.org) for the generous donation of lonafarnib.

## AUTHOR CONTRIBUTIONS

S-I.M., L.B.G., J.D.H. designed the study; S-I.M., N.M., M.W., P.R, Y.K., A.B.R. performed the experiments; S-I.M., N.M., Y.K., J.D.H. analyzed the data; S-I.M., Y.K., D.T.B., G.T., L.B.G., J.D.H. interpreted the results; S-I.M., J.D.H. drafted the MS; all authors read, edited, and approved the final MS.

## SUPPLEMENTAL INFORMATION

## SUPPLEMENTAL FIGURES

**Figure S1.**
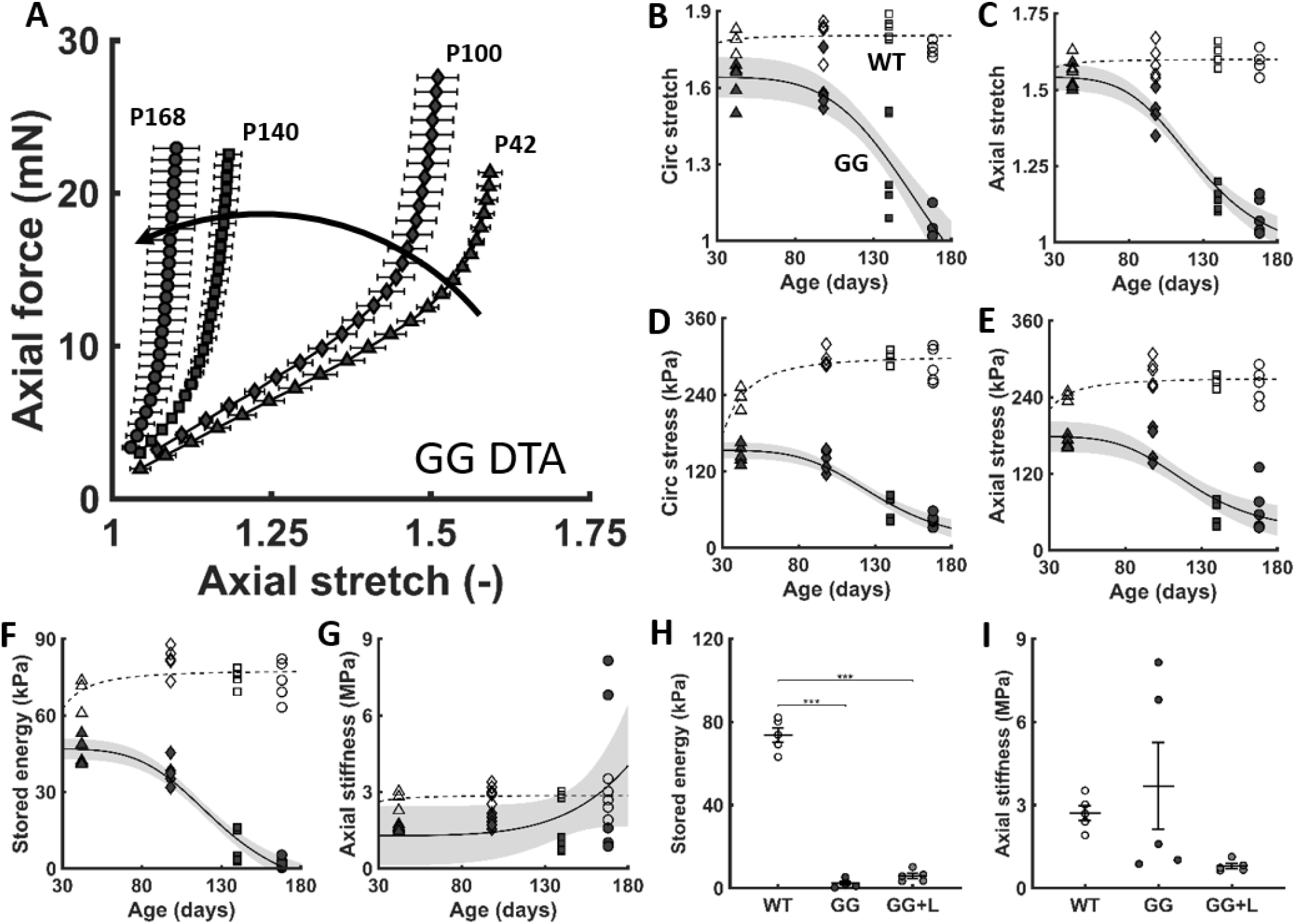
(A) Similar to Figure 1 in the main text, with axial force-stretch responses revealing a progressive structural stiffening of the descending thoracic aorta (DTA) in untreated *Lmna*^*G6096G/G609G*^ progeria (GG) mice as a function of age from post-natal days P42 to P168 (dark curved arrow). (B-G) Multiple geometric and mechanical metrics for the DTA as a function of age, from P42 to P168, for both wild-type (WT – open symbols) and untreated progeria (GG – closed symbols) mice. (H-I) Additional results for WT and both untreated (GG) and lonafarnib-treated (GG+L) aortas, noting that lonafarnib did not improve stored energy or axial stiffness.

**Figure S2.**
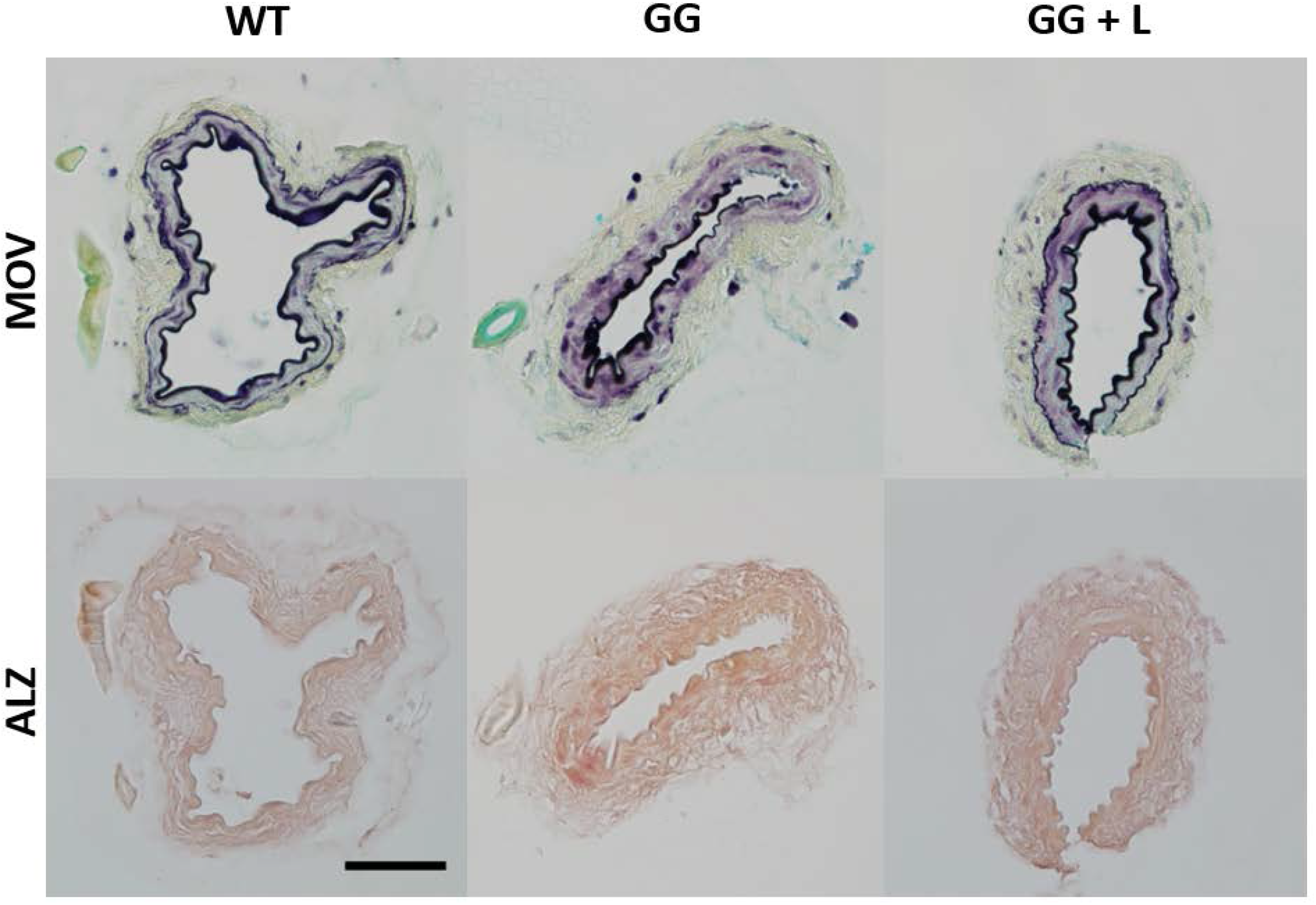
Similar to Figure 4 in the main text, but with standard histology revealing few overt differences in the second branch mesenteric artery (MA) in progeria (GG) relative to wild-type (WT) at post-natal day 168, with little change with lonafarnib treatment (GG+L). First row: Movat-stained sections reveal intact elastic laminae (dark lines), with little accumulation of proteoglycans (blue staining) in progeria. Second row: Alizarin red-stained sections failed to reveal overt calcification in these muscular arteries in progeria. The scale bar is 50 microns.

## SUPPLEMENTAL TABLES

**Table S1.** Tabulated geometric and mechanical metrics for the passive descending thoracic aorta (DTA) from wild-type (WT) as well as untreated (GG), lonafarnib treated (GG+L), and lonafarnib + rapamycin treated (GG+L+R) progeria mice. Pressure-dependent values are evaluated at physiologically relevant, regionally specific values of distending pressure P (in mmHg).

| DTA | WT | GG | GG + L | GG + L + R |
| --- | --- | --- | --- | --- |
|  | n = 5 | n = 5 | n = 5 | n = 5 |
| **Unloaded dimensions** |  |  |  |  |
| Wall thickness ($\mu\text{m}$) | 98 ± 2.4 | 132 ± 6.1 | 125 ± 4.7 | 132 ± 3.6 |
| Outer diameter ($\mu\text{m}$) | 796 ± 8 | 939 ± 21 | 934 ± 36 | 990 ± 15 |
| Axial length (mm) | 4.63 ± 0.15 | 5.28 ± 0.08 | 5.08 ± 0.60 | 4.10 ± 0.26 |
| **Systolic pressure** |  |  |  |  |
|  | P = 126 | P = 101 | P = 101 | P = 101 |
| **Loaded dimensions** |  |  |  |  |
| Outer diameter ($\mu\text{m}$) | 1259 ± 16.0 | 973 ± 20.1 | 1028 ± 21.3 | 987 ± 12.8 |
| Wall thickness ($\mu\text{m}$) | 35 ± 1.3 | 115 ± 7.4 | 95 ± 5.9 | 109 ± 3.2 |
| Inner radius ($\mu\text{m}$) | 594 ± 8.2 | 371 ± 14.5 | 419 ± 8.1 | 385 ± 8.9 |
| *In vivo* axial stretch ($\lambda_z^{\text{iv}}$) | 1.59 ± 0.02 | 1.09 ± 0.03 | 1.14 ± 0.03 | 1.19 ± 0.02 |
| *In vivo* Circumferential Stretch ($\lambda_\theta$) | 1.75 ± 0.01 | 1.07 ± 0.04 | 1.16 ± 0.03 | 1.02 ± 0.01 |
| **Cauchy stresses (kPa)** |  |  |  |  |
| Circumferential, $\sigma_\theta$ | 286 ± 12.2 | 44 ± 4.3 | 60 ± 3.9 | 48 ± 2.6 |
| Axial, $\sigma_z$ | 259 ± 11.6 | 67 ± 17.3 | 52 ± 6.8 | 48 ± 3.3 |
| **Linearized material stiffness (MPa)** |  |  |  |  |
| Circumferential, $\mathcal{E}_{\theta\theta\theta\theta}$ | 1.86 ± 0.11 | 1.71 ± 0.37 | 0.95 ± 0.13 | 0.95 ± 0.13 |
| Axial, $\mathcal{E}_{zzzz}$ | 2.71 ± 0.27 | 3.69 ± 1.57 | 0.80 ± 0.09 | 0.80 ± 0.09 |
| **Stored elastic energy (kPa)** | 74 ± 3.5 | 2 ± 0.9 | 6 ± 1.2 | 3 ± 0.5 |
| **Fixed pressure** |  |  |  |  |
|  | P = 100 | P = 100 | P = 100 | P = 100 |
| **Loaded dimensions** |  |  |  |  |
| Outer diameter ($\mu\text{m}$) | 1192 ± 14.6 | 973 ± 20.1 | 1028 ± 21.4 | 987 ± 12.7 |
| Wall thickness ($\mu\text{m}$) | 37 ± 1.4 | 115 ± 7.4 | 95 ± 5.8 | 109 ± 3.1 |
| Inner radius ($\mu\text{m}$) | 559 ± 7.6 | 371 ± 14.5 | 418 ± 8.1 | 385 ± 8.8 |
| *In vivo* axial stretch ($\lambda_z^{\text{iv}}$) | 1.59 ± 0.02 | 1.09 ± 0.03 | 1.14 ± 0.03 | 1.19 ± 0.02 |
| *In vivo* circumferential stretch ($\lambda_\theta$) | 1.65 ± 0.01 | 1.07 ± 0.04 | 1.16 ± 0.03 | 1.02 ± 0.01 |
| **Cauchy Stresses (kPa)** |  |  |  |  |
| Circumferential, $\sigma_\theta$ | 201 ± 9.0 | 44 ± 4.2 | 59 ± 3.8 | 48 ± 2.6 |
| Axial, $\sigma_z$ | 209 ± 9.1 | 67 ± 17.3 | 52 ± 6.8 | 48 ± 3.3 |
| **Linearized material stiffness (MPa)** |  |  |  |  |
| Circumferential, $\mathcal{E}_{\theta\theta\theta\theta}$ | 1.11 ± 0.07 | 1.69 ± 0.37 | 0.92 ± 0.13 | 0.92 ± 0.13 |
| Axial, $\mathcal{E}_{zzzz}$ | 1.89 ± 0.17 | 3.68 ± 1.57 | 0.80 ± 0.09 | 0.80 ± 0.09 |
| **Stored elastic energy (kPa)** | 60 ± 3.2 | 2 ± 0.9 | 6 ± 1.2 | 3 ± 0.5 |

**Table S2.** Similar to Table S1, but tabulated geometric and mechanical metrics for the descending thoracic aorta (DTA) during vasoactive testing.

|  | DTA | WT | GG | GG + L |
| --- | --- | --- | --- | --- |
|  |  | n = 5 | n = 5 | n = 5 |
| 100 mM KCl | **Loaded configuration** |  |  |  |
| | Relaxed inner radius ($\mu\text{m}$) | 517 ± 14.6 | 372 ± 15.0 | 404 ± 18.4 |
| | Contracted inner radius ($\mu\text{m}$) | 407 ± 8.7 | 371 ± 14.9 | 403 ± 18.4 |
| | Change in inner radius (%) | -21 ± 1.2 | 0 ± 0.2 | 0 ± 0.1 |
|  | **Stress calculations** |  |  |  |
| | Relaxed circumferential stress (kPa) | 142 ± 9.5 | 39 ± 9.5 | 45 ± 4.5 |
| | Contracted circumferential stress (kPa) | 90 ± 5.2 | 39 ± 3.9 | 45 ± 4.4 |
| | Change in circumferential stress (%) | -37 ± 1.8 | -1 ± 0.2 | -1 ± 0.2 |
| 1 $\mu\text{M}$ PE | **Loaded configuration** | | | |
| | Relaxed inner radius ($\mu\text{m}$) | 523 ± 7.7 | 371 ± 15.3 | 405 ± 17.6 |
| | Contracted inner radius ($\mu\text{m}$) | 426 ± 13.6 | 371 ± 15.3 | 404 ± 17.3 |
| | Change in inner radius (%) | -19 ± 2.4 | 0 ± 0.1 | 0 ± 0.1 |
|  | **Stress calculations** |  |  |  |
| | Relaxed circumferential stress (kPa) | 144 ± 7.5 | 39 ± 4.0 | 45 ± 4.5 |
| | Contracted circumferential stress (kPa) | 91 ± 3.8 | 39 ± 3.8 | 45 ± 4.4 |
| | Change in circumferential stress (%) | -36 ± 3.6 | -1 ± 0.2 | -1 ± 0.2 |
| 10 $\mu\text{M}$ Ach + 1 $\mu\text{M}$ PE | **Loaded configuration** | | | |
| | Dilated inner radius ($\mu\text{m}$) | 497 ± 10.5 | 373 ± 15.6 | 406 ± 17.2 |
| | Change in inner radius (%) | 14 ± 1.9 | 1 ± 0.2 | 0 ± 0.2 |
|  | **Stress calculations** |  |  |  |
| | Dilated circumferential stress (kPa) | 131 ± 8.4 | 39 ± 4.0 | 46 ± 4.4 |
| | Change in circumferential stress (%) | 44 ± 7.9 | 2 ± 0.5 | 1 ± 0.2 |

**Table S3.** Tabulated geometric and mechanical metrics for the passive second order mesenteric artery (MA) from wild-type (WT) and both untreated (GG) and lonafarnib treated (GG+L) progeria mice. Pressure-dependent values are evaluated at physiologically relevant, regionally specific values of distending pressure P (in mmHg).

| MA | WT | GG | GG + L |
| --- | --- | --- | --- |
|  | n = 5 | n = 5 | n = 5 |
| **Unloaded dimensions** |  |  |  |
| Wall thickness ($\mu\text{m}$) | 50 ± 1.7 | 55 ± 5.2 | 51 ± 2.5 |
| Outer diameter ($\mu\text{m}$) | 225 ± 8 | 225 ± 10 | 167 ± 12 |
| Axial length (mm) | 2.65 ± 0.28 | 3.44 ± 0.52 | 4.94 ± 0.27 |
| **Systolic pressure** |  |  |  |
|  | P = 76 | P = 61 | P = 61 |
| **Loaded dimensions** |  |  |  |
| Outer diameter ($\mu\text{m}$) | 282 ± 11.3 | 234 ± 8.3 | 184 ± 18.7 |
| Wall thickness ($\mu\text{m}$) | 19 ± 0.9 | 35 ± 4.0 | 27 ± 3.5 |
| Inner radius ($\mu\text{m}$) | 122 ± 5.4 | 82 ± 2.0 | 65 ± 9.9 |
| *In vivo* axial stretch ($\lambda_z^{\text{iv}}$) | 1.75 ± 0.04 | 1.36 ± 0.10 | 1.47 ± 0.07 |
| *In vivo* Circumferential Stretch ($\lambda_\theta$) | 1.51 ± 0.03 | 1.18 ± 0.04 | 1.35 ± 0.09 |
| **Cauchy stresses (kPa)** |  |  |  |
| Circumferential, $\sigma_\theta$ | 65 ± 3.7 | 20 ± 3.0 | 22 ± 4.5 |
| Axial, $\sigma_z$ | 109 ± 8.9 | 33 ± 10.2 | 44 ± 17.0 |
| **Linearized material stiffness (MPa)** |  |  |  |
| Circumferential, $\mathcal{E}_{\theta\theta\theta\theta}$ | 0.76 ± 0.09 | 0.24 ± 0.04 | 0.14 ± 0.04 |
| Axial, $\mathcal{E}_{zzzz}$ | 1.85 ± 0.26 | 0.75 ± 0.22 | 0.55 ± 0.23 |
| **Stored elastic energy (kPa)** | 19 ± 1.6 | 4 ± 1.7 | 7 ± 2.1 |
| **Fixed pressure** |  |  |  |
|  | P = 60 | P = 60 | P = 60 |
| **Loaded dimensions** |  |  |  |
| Outer diameter ($\mu\text{m}$) | 275 ± 11.3 | 234 ± 8.3 | 184 ± 18.7 |
| Wall thickness ($\mu\text{m}$) | 20 ± 0.9 | 35 ± 4.0 | 27 ± 3.5 |
| Inner radius ($\mu\text{m}$) | 118 ± 5.5 | 82 ± 1.9 | 65 ± 9.9 |
| *In vivo* axial stretch ($\lambda_z^{\text{iv}}$) | 1.75 ± 0.04 | 1.36 ± 0.10 | 1.47 ± 0.07 |
| *In vivo* circumferential stretch ($\lambda_\theta$) | 1.46 ± 0.03 | 1.18 ± 0.04 | 1.35 ± 0.09 |
| **Cauchy Stresses (kPa)** |  |  |  |
| Circumferential, $\sigma_\theta$ | 48 ± 2.9 | 20 ± 3.0 | 21 ± 4.4 |
| Axial, $\sigma_z$ | 93 ± 8.4 | 33 ± 10.2 | 44 ± 17.0 |
| **Linearized material stiffness (MPa)** |  |  |  |
| Circumferential, $\mathcal{E}_{\theta\theta\theta\theta}$ | 0.47 ± 0.06 | 0.23 ± 0.04 | 0.14 ± 0.04 |
| Axial, $\mathcal{E}_{zzzz}$ | 1.33 ± 0.16 | 0.74 ± 0.22 | 0.54 ± 0.23 |
| **Stored elastic energy (kPa)** | 17 ± 1.5 | 4 ± 1.7 | 7 ± 2.1 |

**Table S4.** Tabulated geometric and mechanical metrics for the second order mesenteric artery (MA), during vasoactive testing, from wild-type (WT) and both untreated (GG) and lonafarnib treated (GG+L) progeria mice.

| MA |  | WT | GG | GG + L |
| --- | --- | --- | --- | --- |
|  |  | n = 5 | n = 4 | n = 5 |
| 100 mM KCl | **Loaded configuration** |  |  |  |
| | Relaxed inner radius ($\mu\text{m}$) | 122 ± 2.8 | 84 ± 3.1 | 83 ± 4.1 |
| | Contracted inner radius ($\mu\text{m}$) | 54 ± 5.7 | 64 ± 13.2 | 47 ± 4.7 |
| | Change in inner radius (%) | -56 ± 4.7 | -23 ± 15.6 | -43 ± 5.4 |
|  | **Stress calculations** |  |  |  |
| | Relaxed circumferential stress (kPa) | 49 ± 4.5 | 21 ± 4.5 | 34 ± 3.5 |
| | Contracted circumferential stress (kPa) | 25 ± 3.1 | 17 ± 3.3 | 21 ± 0.6 |
| | Change in circumferential stress (%) | -37 ± 20.5 | -36 ± 16.2 | -46 ± 19.1 |
| 1 $\mu\text{M}$ PE | **Loaded configuration** | | | |
| | Relaxed inner radius ($\mu\text{m}$) | 121 ± 2.4 | 85 ± 4.2 | 69 ± 5.6 |
| | Contracted inner radius ($\mu\text{m}$) | 73 ± 1.8 | 75 ± 8.4 | 41 ± 5.7 |
| | Change in inner radius (%) | -40 ± 0.7 | -11 ± 8.5 | -41 ± 4.4 |
|  | **Stress calculations** |  |  |  |
| | Relaxed circumferential stress (kPa) | 49 ± 4.6 | 22 ± 0.8 | 26 ± 3.0 |
| | Contracted circumferential stress (kPa) | 33 ± 2.6 | 20 ± 2.0 | 17 ± 1.6 |
| | Change in circumferential stress (%) | -16 ± 2.0 | -2 ± 1.4 | -9 ± 1.8 |
| 10 $\mu\text{M}$ Ach + 1 $\mu\text{M}$ PE | **Loaded configuration** | | | |
| | Dilated inner radius ($\mu\text{m}$) | 120 ± 1.7 | 79 ± 6.5 | 66 ± 7.6 |
| | Change in inner radius (%) | 64 ± 3.8 | 7 ± 3.7 | 62 ± 15.0 |
|  | **Stress calculations** |  |  |  |
| | Dilated circumferential stress (kPa) | 49 ± 5.4 | 20 ± 2.1 | 24 ± 2.6 |
| | Change in circumferential stress (%) | 48 ± 5.5 | 1 ± 2.3 | 37 ± 12.9 |

